# Dissection and validation of minor quantitative trait loci (QTLs) conferring grain size and weight in rice

**DOI:** 10.1101/511139

**Authors:** Ping Sun, Yuanyuan Zheng, Pingbo Li, Hong Ye, Hao Zhou, Guanjun Gao, Yuqing He

## Abstract

Grain size and weight contribute greatly to the grain yield of rice. In order to identify minor QTLs conferring grain size and weight, an F_2_ population derived from a cross between two *indica* rice lines showing small difference on grain size, Guangzhan 63-4S (GZ63-4S) and Dodda, and its derived F_2:3_ population were developed and used for QTL analysis. Totally, 36 QTLs for grain size and weight were detected, and 7 were repeatedly detected, of which the number of beneficial alleles was contributed roughly equally by the two parents. In order to further validate effects of QTLs detected, a BC_1_F_2_ population derived from a backcross of a mixture of F_2_ lines with GZ63-4S was developed and subjected to QTL selection. Heterozygous regions of 3 QTLs, *qGS3*, *qTGW6.2* and *qGT7* were identified, and corresponding near-isogenic lines (NILs) of each QTL were constructed with three rounds of self-crosses. In the background of NILs, *qGS3* was responsible for GL, LWR, GT and TGW, *qTGW6.2* was for GL and TGW, and *qGT7* was for GT and TGW. These results have laid the foundation of further fine mapping and cloning of underlying genes, and could be of great use in breeding and improvement of rice lines with desirable size and yield.

## Introduction

Rice is one of the staple crops worldwide, and feeds more than half of the world’s population. In the face of continuously increasing population and reduced arable land, how to further improve the grain yield of rice is a major concern of scientists and breeders. Grain size, characterized by four factors viz., grain length (GL), grain width (GW), length-to-width ratio (LWR) and grain thickness (GT), contributes greatly to grain weight, which is a key determinant of grain yield [1]. Therefore, dissection of the genetic basis that underlies grain size and weight would be of great use in developing rice lines with high grain yield.

Considerable efforts have been made to investigate the genetic basis of grain size and weight in the past two decades, and results showed that the four factors of grain size, GL, GW, LWR and GT, and thousand-grain weight (TGW) are quantitative traits, and subjected to control of many genes [2, 3]. Up to now, large numbers of quantitative trait loci (QTLs) have been identified, however, only a small proportion of QTLs displaying large effect have been cloned, such as *GS3* [4, 5], *OsMADS1* [6, 7], *GL3.3*/*TGW3* [8–10], *GW5*/*GSE5* [11, 12], *GS5* [13], *GW8* [14], *GS2*/*GL2* [15–17], *GL7*/*GW7* [18, 19], etc. Although the knowledge of molecular regulation of grain size and weight has greatly increased, the mining and cloning of more QTLs, especially minor QTLs, is still of great importance to have a better understanding of underlying mechanisms and provide breeding programs with valuable gene resources.

Rice lines displaying large difference on grain size and weight were always selected to develop segregating populations for QTL analysis, which resulted in the repeated detection of several major QTLs/genes. For example, two major genes for grain size, *GW2* and *GL3.1* were identified and cloned from genetic populations derived from FAZ1 and WY3, of which the TGW values differ by 23.12g [20, 21]. The two genes above, together with another two major genes, *GS3* and *GW5*/*GSE5*, contributed to the huge variation of grain size and weight between N411 and N643, of which the TGW values differ by 54.33g [22]. The existence of major genes is likely to interfere the mapping and validation of minor QTLs, exemplified by the fine mapping of *GS5* [13]. Therefore, in order to identify minor QTLs for grain size and weight, rice lines displaying small difference should be preferred.

Quantitative traits are easily affected by environment, which leads to the instability of QTL detection. Therefore, genetic validation of QTLs is of great necessity in further breeding utilization or cloning. The most frequently used method is evaluation the effect of a QTL using near-isogenic lines (NILs), which are lines that carry segregating regions at target QTL but homozygous regions in the rest of genome [23]. NILs for a QTL are always developed by backcrossing lines carrying the QTL region from donor to the receipt several times until the non-target QTL regions were completely from the receipt, which could achieve the simultaneous improvement of target traits of recipient [24, 25]. Another simple method is to select lines carrying segregating target QTL regions from inbred populations that have undertaken several rounds of self-crosses, also known as residual heterozygous lines (RHLs) [26, 27]. This method is sometimes utilized for absence of laborious hybridization work. The NIL of *Ghd8*, a major QTL with pleiotropic effects on grain yield, heading date and plant height, was constructed by screening lines carrying segregation target regions from a RIL population of the F_7_ generation [28, 29].

In this study, in order to identify minor QTLs for grain size, two *indica* rice lines displaying small difference, Guangzhan 63-4S (GZ63-4S) and Dodda were selected to develop the F_2_ and derived F_2:3_ populations, and QTL analysis of grain size and weight were performed. In order to validate QTL detected, lines carrying heterozygous QTL regions were screened from a BC_1_F_2_ population derived from a backcross of a mixture of F_2_ lines with GZ63-4S. NILs of three QTLs were developed by a series of self-crosses of screened BC_1_F_2_ lines, and further used for evaluation their genetic effect on grain size and TGW.

## Materials and methods

### Population development and cultivation

Guangzhan 63-4S (GZ63-4S) is a leading *indica* two-line male sterile line developed by the China North Japonica Hybrid Rice Research Center and Hefei Fengle Seed Company, and has been mated with many restorer lines to produce promising hybrid combinations in recent years [30]. Dodda is an *indica* cultivar with unknown origin, belonging to the core germplasm collections of our lab. The TGW values of GZ63-4S and Dodda differ by less than 10 g (data not shown).

As displayed in Fig.1, 1000 F_2_ lines were produced from a cross between GZ63-4S and Dodda, and were subjected to selection of the *TMS5* locus conditioning thermo-sensitive genic male sterility with a closely linked marker [31]. 214 lines carrying homozygous *TMS5* regions were selected to make up the F_2_ mapping population, which was further self-crossed to produce the F_2:3_ population. Both the F_2_ and F_2:3_ population was exploited to map QTLs for grain size and TGW. In addition, 1200 BC_1_F_2_ lines were produced by backcrossing a mixture of F_2_ lines to GZ63-4S, followed by a self-cross. These lines were subjected to *TMS5* selection, and 250 lines carrying homozygous *TMS5* regions were selected to perform heterozygous QTL regions screening with flanking makers in the mapping process (Table 2). BC_1_F_2_ lines carrying heterozygous QTL regions were further self-crossed three times to produce the BC_1_F_5_ populations, which were utilized to validate the effect of QTLs. The F_2_, F_2:3_ and BC_1_F_5_ populations were planted in year 2014, 2015 and 2018, respectively, during the normal rice growing seasons at the Experimental Farm of Huangzhong Agricultural University in Wuhan, China. Each F_3_ line consisted of 12 plants, and each BC_1_F_5_ population consisted of 100 plants.

**Fig. 1.**
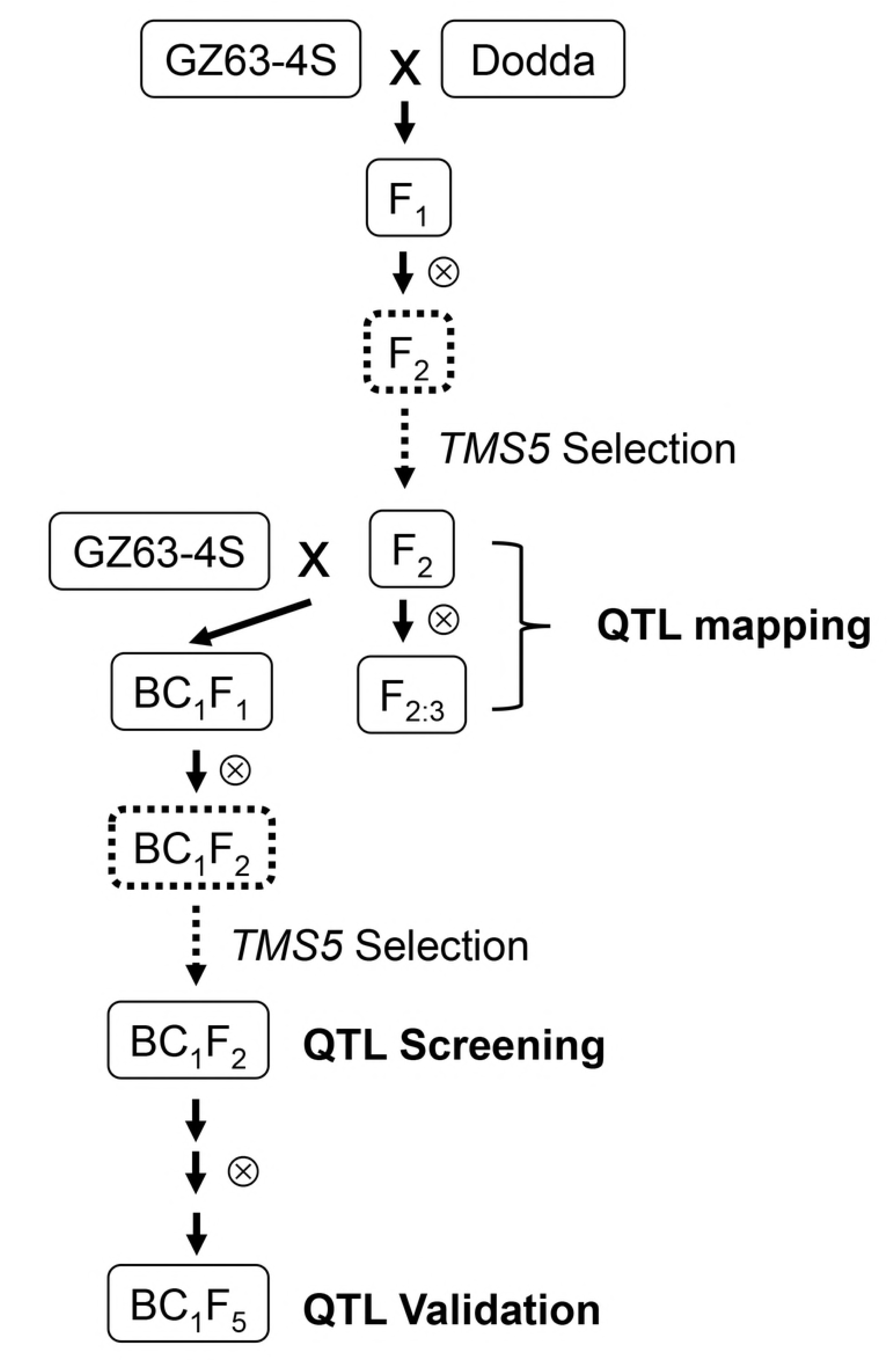
Schematic representation of the experimental design.

**Table 1.** Correlations of GL, GW, LWR, GT and TGW of the F_2_ and F_2:3_ populations in year 2014 and 2015

|  | GL14 | GL15 | GW14 | GW15 | LWR14 | LWR15 | GT14 | GT15 | TGW14 | TGW15 |
| --- | --- | --- | --- | --- | --- | --- | --- | --- | --- | --- |
| GL14 |  |  |  |  |  |  |  |  |  |  |
| GL15 | 0.731** |  |  |  |  |  |  |  |  |  |
| GW14 | 0.157 | 0.055 |  |  |  |  |  |  |  |  |
| GW15 | 0.046 | 0.139 | 0.489** |  |  |  |  |  |  |  |
| LWR14 | 0.525** | 0.453** | -0.728** | -0.424** |  |  |  |  |  |  |
| LWR15 | 0.477** | 0.585** | -0.348** | -0.692** | 0.657** |  |  |  |  |  |
| GT14 | 0.387** | 0.381** | 0.126 | 0.269** | 0.144 | 0.053 |  |  |  |  |
| GT15 | 0.225* | 0.452** | 0.109 | 0.235* | 0.057 | 0.118 | 0.677** |  |  |  |
| TGW14 | 0.656** | 0.558** | 0.378** | 0.415** | 0.129 | 0.046 | 0.739** | 0.495** |  |  |
| TGW15 | 0.484** | 0.712** | 0.261** | 0.450** | 0.093 | 0.132 | 0.584** | 0.707** | 0.700** |  |

GL14, GL in 2014; GL15, GL in 2015; GW14, GW in 2014; GW15, GW in 2015; LWR14, LWR in 2014; LWR15, LWR in 2015; GT14, GT in 2014; GT15, GT in 2015; TGW14, TGW in 2014; TGW15, TGW in 2015.
\*, \*\* Significantly at $P<0.05$ and $P<0.01$ , respectively.

**Table 2.**
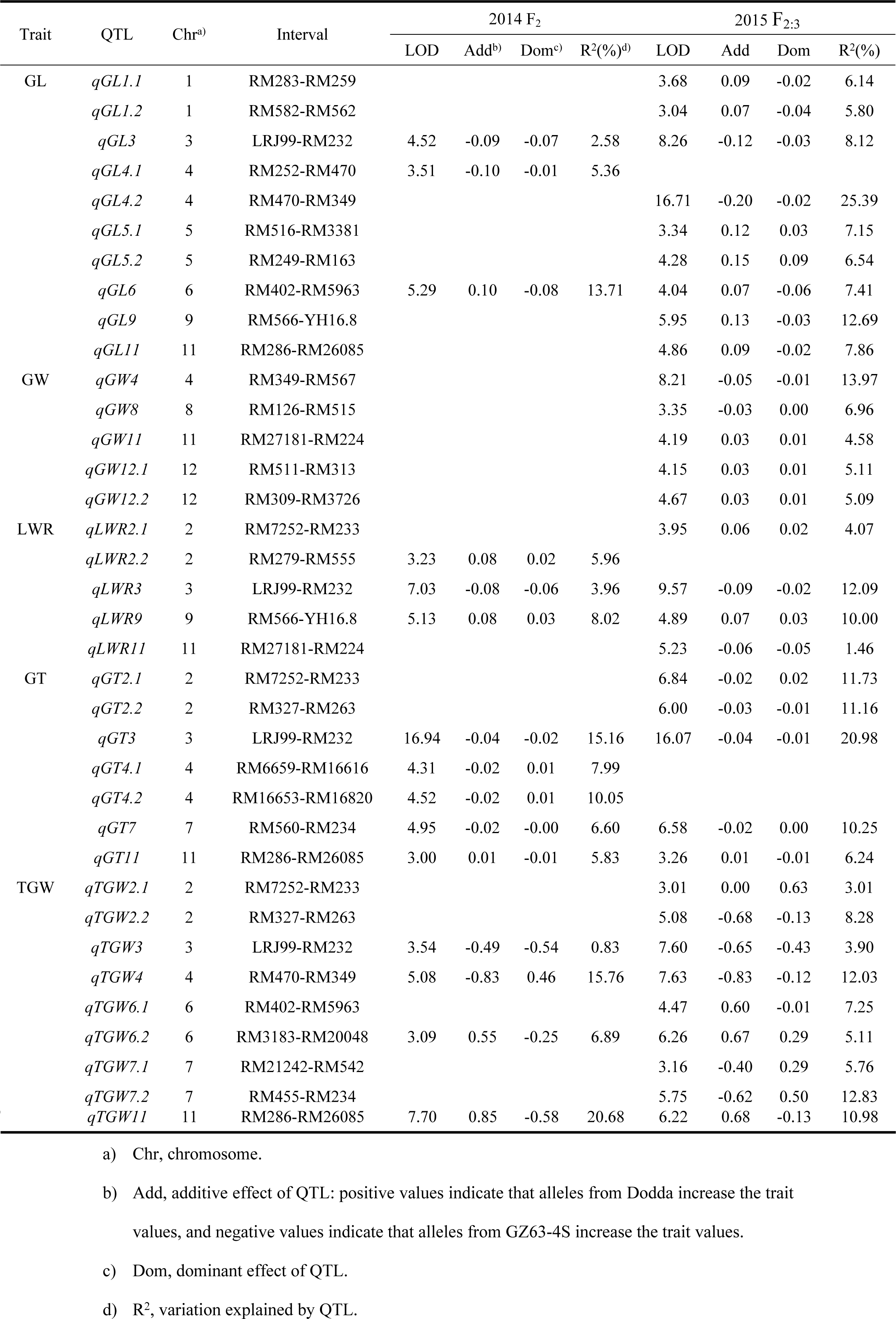
QTLs detected for GL, GW, LWR, GT and TGW in the F_2_ and F_2:3_ populations in year 2014 and 2015, respectively

### Trait evaluation

GL, GW, LWR and TGW were measured with more than 200 grains per line or plant using the yield traits scorer [32]. GT was determined for each grain individually using an electronic digital caliper (Guanglu Measuring Instrument Co. Ltd., China), and thirty grain values were averaged for each line or plant. For the F_2:3_ population, the phenotypic value of each line was the average value of 12 plants.

### Genetic map construction

A total of more than 1000 simple sequence repeat markers or insert/deletion markers were employed to screen for polymorphic markers between GZ63-4S and Dodda, and 143 markers were identified. Among that, 111 markers were selected to perform genotyping of the F_2_ population with 4% polyacrylamide gels migration and silver staining [33]. A genetic linkage map was constructed using MapMaker/Exp3.0 with the Kosambi mapping function [34].

### Data analysis

Correlation analysis was performed using the data analysis module in Microsoft Office Excel 2016. QTL analysis was performed by composite interval mapping using the software package QTLCartographer V2.5 with a logarithm of odds (LOD) threshold of 3.0 [35]. ANNOVA analysis was performed using the IBM SPSS Statistics 22.

## Results

### Phenotypic variation and correlation of the F_2_ and F_2:3_ populations

GZ63-4S is a typical photoperiod- and thermo-sensitive genic male sterile line, and shows male sterility in the normal growing seasons in Wuhan. Therefore, the seeds could not be harvested, which abolished comparison of grain size and TGW between the two parents. All the five traits of the F_2_ and F_2:3_ populations showed continuous variation and followed normal distribution in year 2014 and 2015, respectively (Fig. 2).

**Fig. 2.**
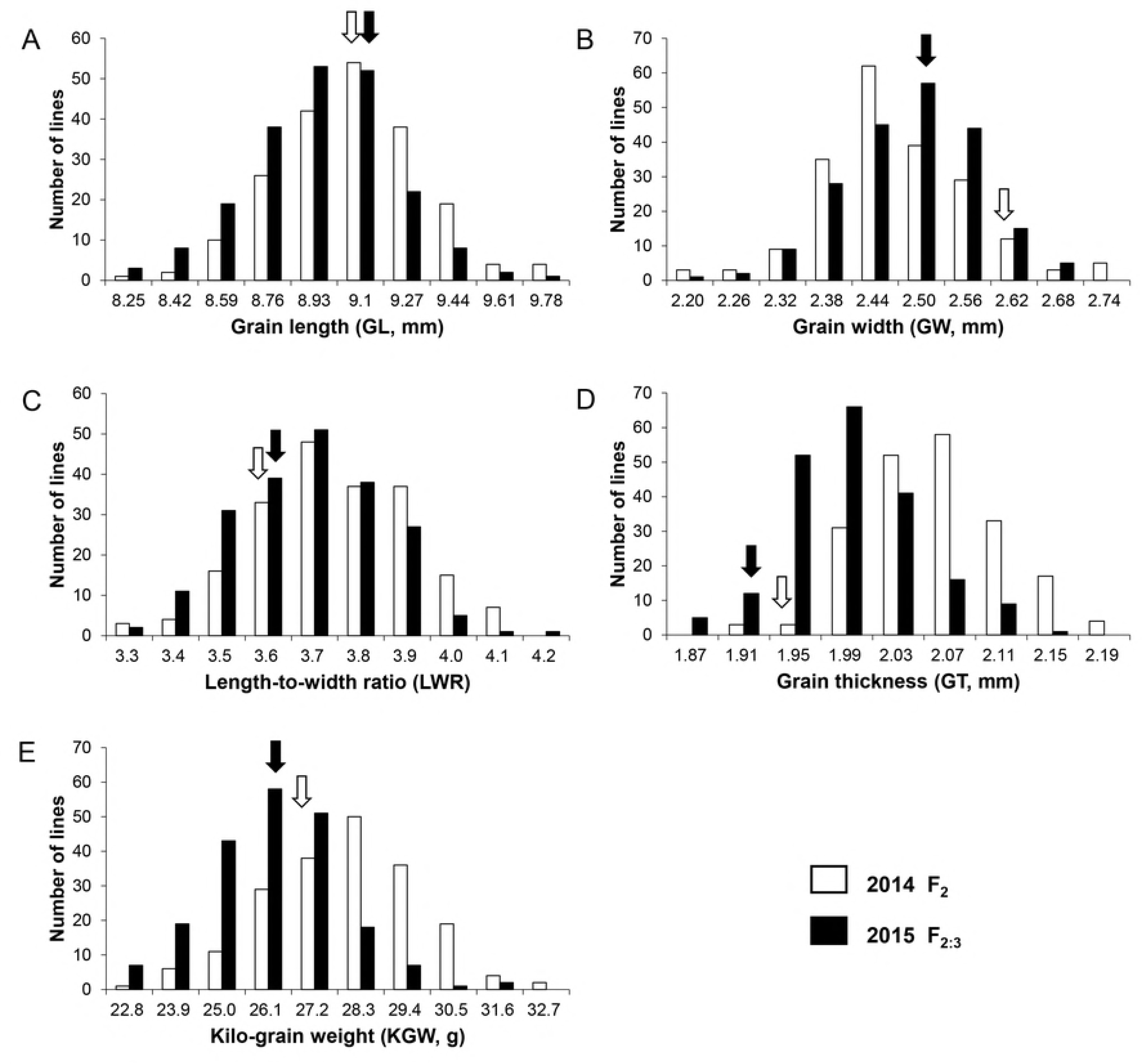
Frequency distribution of the F_2_ and F_2:3_ populations for GL, GW, LWR, GT and TGW in year 2014 and 2015. Arrow indicates the value of Dodda.

All the four grain size factors were significantly positively correlated with TWG in both years, except for LWR (Table 1). GL was significantly positively correlated with LWR and GT in both years, while GW was only significantly negatively correlated with LWR in both years. The three highest correlation coefficients were observed between GW and TGW in year 2014, GL in two years, and GW and LWR in year 2014, with values of 0.739, 0.731 and 0.728, respectively.

### QTLs detected in the F_2_ and F_2:3_ populations

#### GL

Ten QTLs for GL were detected in two populations and distributed on seven chromosomes, with phenotypic variation explained by each QTL ranging from 2.58% to 25.39% (Table 2, Fig.3). Among those, the beneficial alleles of *qGL3*, *qGL4.1* and *qGL4.2* were from GZ63-4S, while that of others were from Dodda. Two QTLs, *qGL3* and *qGL6* were repeatedly detected, and explained 2.58% and 13.71% of the variation in the F_2_ population, and 8.12% and 7.41% of the variation in the F_2:3_ population, respectively. The remaining QTLs were detected only in the F_2:3_ population, excluding *qGL4.1*.

**Fig. 3.**
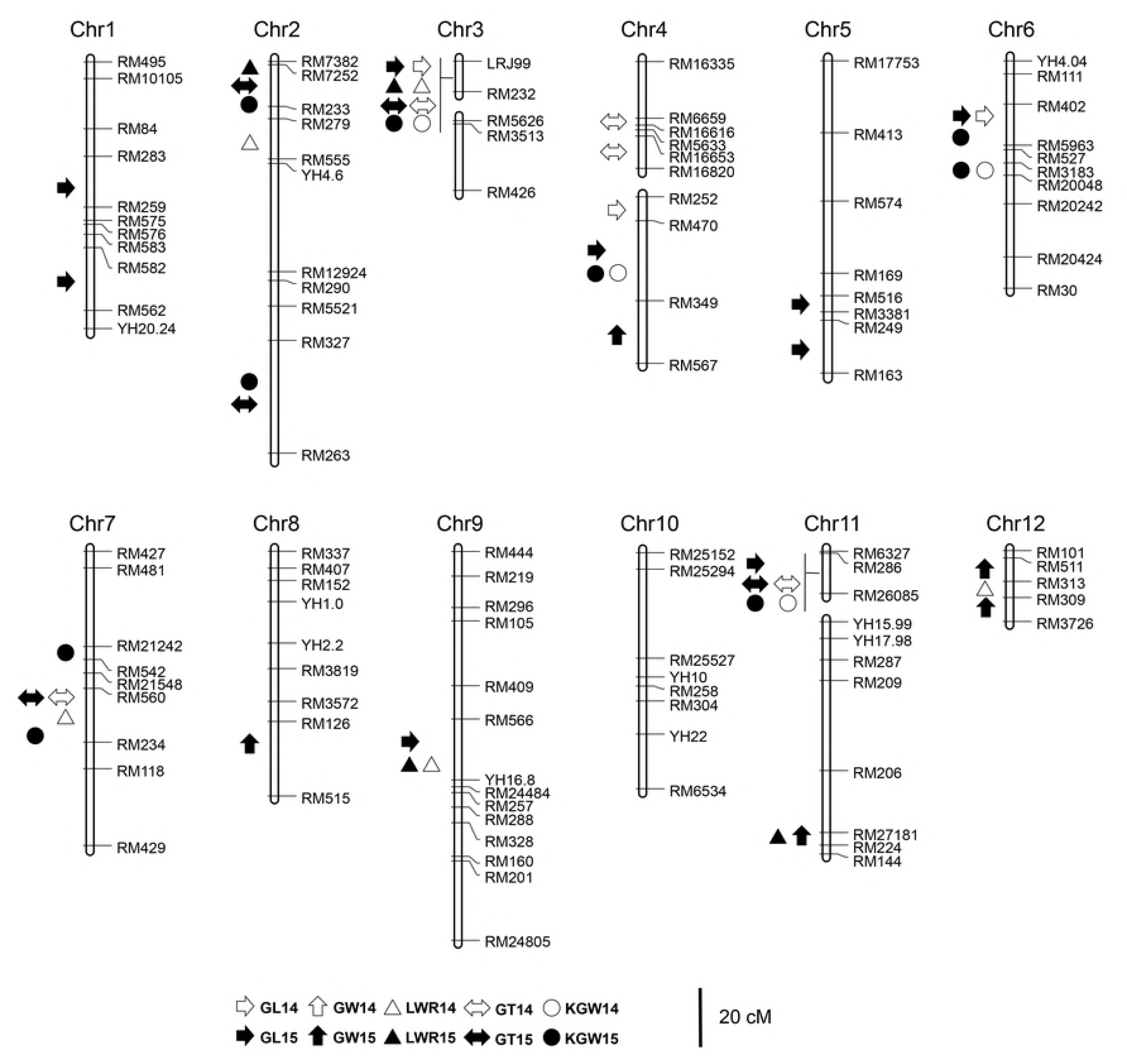
Distribution of putative QTLs for GL, GW, LWR, GT and TGW identified in the F_2_ and F_2:3_ populations on the linkage map. GL14, QTLs for GL detected in the F_2_ population in year 2014; GL15, QTLs for GL detected in the F_2:3_ population in year 2014. The QTLs for GW, LWR, GT and TGW are represented as the same manner as that for GL.

#### GW

Five QTLs were detected for GW in the F_2:3_ population, while none in the F_2_ population (Table 2, Fig.3). Among those, the beneficial alleles of two were from GZ63-4S, while that of the other three were from Dodda.

#### LWR

Five QTLs for LWR were identified in the two populations, and distributed on four chromosomes (Table 2, Fig.3). Among those, two QTLs, *qLWR3* and *qLWR9*, were repeatedly detected, and displayed nearly the same values of additive effect in opposite direction. The remaining were minor QTLs accounting for less than 6% of the variation and were detected only in one population.

#### GT

Seven QTLs were identified for GT in the two populations and were distributed on chromosome 2, 3, 4, 7 and 11 (Table 2, Fig.3). The beneficial allele of all eight QTLs were from GZ63-4S, except for that of *qGT11*. The three QTLs, *qGT3*, *qGT7* and *qGT11* were stably detected, and explained 15.16%, 6.60% and 5.83% of the variation in the F_2_ population, and 20.98%, 10.25% and 6.24% of the variation in the F_2:3_ population, respectively. *qGT2.1* and *qGT2.2* were only detected in the F_2:3_ population, while *qGT7* and *qGT11* were only in the F_2_ population.

#### TGW

Nine QTLs for TGW were detected in the two populations, which were distributed on chromosome 2, 3, 4, 6, 7 and 11 (Table 2, Fig.3). Among those, four QTLs, *qTGW3*, *qTGW4*, *qTGW6.2* and *qTGW11*, were repeatedly detected, which accounted for 0.83%, 15.76%, 6.89% and 20.68% of the variation in the F_2_ population, and 3.90%, 12.03%, 5.11% and 10.98% of the variation in the F_2:3_ population, respectively. The beneficial alleles of *qTGW3* and *qTGW4* were from GZ63-4S, while that of *qTGW6.2* and *qTGW11* were from Dodda. The remaining QTLs were all only detected in the F_2:3_ population. The region flanked by marker LRJ99 and RM232 on chromosome 3 and consisting of four QTLs, was responsible for GL, LWR, GT and TGW in both the F_2_ and F_2:3_ population, and was term *qGS3*, hereafter.

### Validation of QTL effects using NILs

In the BC_1_F_2_ population derived from backcrossing some F_2_ lines with GZ63-4S, heterozygous regions were screened for QTLs repeatedly detected in both the F_2_ and F_2:3_ populations or QTLs accounting for more that 10% of variation in one population using flanked markers (Fig.2, Table 1). Lines carrying heterozygous regions of three QTLs, *qGS3*, *qTGW6.2* and *qGT7*, were identified respectively, and were self-crossed three times to produce NIL populations for each QTL.

In the NIL population of *qGS3*, significant differences were observed in the average values of GL, GT and TGW among the three genotypes, *qGS3*^*Dodda*^, *qGS3*^*H*^ and *qGS3*^*GZ63-4S*^ (Table 3). Compared to NIL (*qGS3*^*Dodda*^), NIL (*qGS3*^*GZ63-4S*^) showed increased values by 0.21 mm in GL, 0.10 in LWR, 0.07mm in GT and 1.47g in TGW. For *qTGW6.2*, significant differences were observed in the average values of GL and TGW between *qTGW6.2*^*Dodda*^ and *qTGW6.2*^*GZ63-4S*^, while no difference between *qTGW6.2*^*H*^ and *qTGW6.2*^*GZ63-4S*^ in the NIL population (Table 4). Compared to NIL (*qTGW6.2*^*Dodda*^), NIL (*qTGW6.2*^*GZ63-4S*^) showed decreased values by 0.14 mm in GL and 1.24g in TGW.

**Table 3.** Genetic effect of *qGS3* in the NIL population

|  | Number | GL (mm) | GW (mm) | LWR | GT (mm) | TGW (g) |
| --- | --- | --- | --- | --- | --- | --- |
|  | of lines | Mean±SE | Mean±SE | Mean±SE | Mean±SE | Mean±SE |
| NIL(*qGS3*<sup>Dodda</sup>) | 21 | 9.06±0.14 a | 2.77±0.05 | 3.31±0.08 a | 1.97±0.02 a | 24.10±0.96 a |
| NIL(*qGS3*<sup>H</sup>) | 40 | 9.15±0.14 b | 2.77±0.06 | 3.33±0.07 a | 2.00±0.02 b | 24.88±1.10 b |
| NIL(*qGS3*<sup>GZ63-4S</sup>) | 19 | 9.27±0.11 c | 2.75±0.01 | 3.41±0.05 b | 2.04±0.03 c | 25.57±1.08 c |

Lower-case letters indicate statistically significant ( $P<0.05$ ) differences between the mean values within each column (Duncan test).
NIL(*qGS3*<sup>Dodda</sup>) and NIL(*qGS3*<sup>GZ63-4S</sup>) are the lines carrying homozygous *qGS3* regions from Dodda and GZ63-4S in the NIL population, respectively, while NIL(*qGS3*<sup>H</sup>) is the line carrying heterozygous *qGS3* regions.

**Table 4.** Genetic effect of *qTGW6.2* in the NIL population

|  | Number | GL (mm) | GW (mm) | LWR | GT (mm) | TGW (g) |
| --- | --- | --- | --- | --- | --- | --- |
|  | of lines | Mean±SE | Mean±SE | Mean±SE | Mean±SE | Mean±SE |
| NIL(*qTGW6.2*<sup>Dodda</sup>) | 19 | 9.64±0.11 b | 2.64±0.05 b | 3.70±0.06 | 2.15±0.03 | 26.86±1.25 b |
| NIL(*qTGW6.2*<sup>H</sup>) | 34 | 9.55±0.11 a | 2.61±0.06 a | 3.71±0.08 | 2.14±0.03 | 25.96±1.10 a |
| NIL(*qTGW6.2*<sup>GZ63-4S</sup>) | 18 | 9.50±0.11 a | 2.61±0.04 ab | 3.69±0.07 | 2.14±0.03 | 25.62±1.08 a |

Lower-case letters indicate statistically significant ( $P<0.05$ ) differences between the mean values within each column (Duncan test).
NIL(*qTGW6.2*<sup>Dodda</sup>) and NIL(*qTGW6.2*<sup>GZ63-4S</sup>) are the lines carrying homozygous *qTGW6.2* regions from Dodda and GZ63-4S in the NIL population, respectively, while NIL(*qTGW6.2*<sup>H</sup>) is the line carrying heterozygous *qTGW6.2* regions.

For *qGT7*, significant differences were observed in the average values of GT among the three genotypes, *qGT7*^*Dodda*^, *qGT7*^*H*^ and *qGT7*^*GZ63-4S*^, and in that of TGW between *qGT7*^*Dodda*^ and *qGT7*^*GZ63-4S*^ in the NIL population (Table 5). Compared to NIL (*qGT7*^*Dodda*^), NIL (*qGT7*^*GZ63-4S*^) showed increased values by 0.07 mm in GT and 1.27g in TGW.

**Table 5.** Genetic effect of *qGT7* in the NIL population

|  | Number | GL (mm) | GW (mm) | LWR | GT (mm) | TGW (g) |
| --- | --- | --- | --- | --- | --- | --- |
|  | of lines | Mean±SE | Mean±SE | Mean±SE | Mean±SE | Mean±SE |
| NIL(*qGT7<sup>Dodda</sup>*) | 14 | 9.66±0.12 | 2.57±0.06 | 3.81±0.10 b | 1.96±0.03 a | 21.84±1.20 a |
| NIL(*qGT7<sup>H</sup>*) | 21 | 9.62±0.14 | 2.61±0.08 | 3.74±0.08 a | 1.99±0.03 b | 22.81±1.19 b |
| NIL(*qGT7<sup>GZ63-4S</sup>*) | 13 | 9.60±0.13 | 2.60±0.06 | 3.76±0.07 ab | 2.03±0.03 c | 23.17±1.05 b |

Lower-case letters indicate statistically significant ( $P<0.05$ ) differences between the mean values within each column (Duncan test).
NIL(*qGT7<sup>Dodda</sup>*) and NIL(*qGT7<sup>GZ63-4S</sup>*) are the lines carrying homozygous *qGT7* regions from Dodda and GZ63-4S in the NIL population, respectively, while NIL(*qGT7<sup>H</sup>*) is the line carrying heterozygous *qGT7* regions.

## Discussion

### Evaluation of grain size and weight

Genotyping and phenotyping are two key processes in genetic analysis. With the completion of high-quality genome sequences of several rice cultivars and development of sequencing techniques, genotyping a population is becoming increasingly simple and cheap [36, 37]. Therefore, high-throughput and time-saving methods of phenotyping are in urgent need. In previous studies, grain size was always evaluated using electronic digital-display vernier caliper, and about 30 randomly chosen filled grains was used for each line, which is both pains taking and time consuming [19, 38]. *SmartGrain* is a phenotyping software developed for measuring grain size through image analysis, which improved greatly the throughput, but is still time consuming for the separation of adjacent seeds in the scanning process [39]. In this study, evaluation of GL, GW, LWR, and TGW was performed using the yield traits scorer (YTS) that could fulfil the measurement of a rice line represented by about 500 seeds within one minute [28]. Therefore, the YTS dramatically increases the amount of seeds evaluated and reduces the time of phenotyping, demonstrating its great power in phenotype evaluation.

### Minor QTLs for grain size and weight

In this study, a total of 37 QTLs were identified for GL, GW, LWR, GT and TGW in the F_2_ and F_2:3_ populations, and 7 QTL regions were repeatedly detected, of which the additive effects were far less than that of cloned major genes for grain size, such as *GS3*, *GL3.1/qGL3*, *GW5*/*GSE5* and *GW2* [4, 5, 11, 12, 20-22], demonstrating minor QTLs for grain size and weight. Moreover, the number of beneficial alleles was contributed roughly equally by the two parents, indicating that novel minor QTLs could be detected from rice lines that differ little in grain size.

Among QTLs detected, *qGS3*, the pleiotropic QTL for GL, LWR, GT and TGW on chromosome 3, is co-located with *OsMADS1*, of which a natural allele was reported responsible for GL, GT and TGW in two separate studies [6, 7]. However, the difference of GL between the two NILs in Yu et al. (2018) was almost twice that between our NILs, while the difference of GT was the same [7]. An appropriate explanation is that another gene for GL in the Nipponbare background may interact with *OsMADS1* to amplify the difference in the NILs, as reported by Xia et al. (2018) [9]. Therefore, *qGS3* is likely to be *OsMADS1*. In addition, the region of *qTGW6.2* overlaps with that of two QTLs for TGW in the chromosomal segment substitution lines (CSSLs) population derived from Yamadanishiki or Takanari in the background of Koshihikari, respectively [40, 41]. *qGT7*, the QTL for GT on chromosome 7, is co-located with a region for GL, GW, LWR and GT reported by Liu et al (2015), which contains *GL7*/*GW7*, a major gene influencing GL and GW simultaneously [18, 19, 42]. As *qGT7* has no effect on GL and GW in the NIL background, it is a novel gene different from *GL7*/*GW7*. The region of *qLWR9* was also detected for LWR only by Yin et al. (2015) [43]. The remaining QTLs are seldom reported, or maybe novel.

### Validation of minor QTLs using NILs

QTLs detected in primary populations are sometimes unstable, and thus should be further validated, especially for minor QTLs. The best way to validate QTLs is the use of NILs. In this study, lines carrying heterozygous QTL regions were screened from the BC_1_F_2_ population, in case of the loss of target regions in subsequent self-crosses. Then, selected lines were subjected to three rounds of self-crosses, in order to reduce the heterozygosity of non-target regions. The method we preferred ensures that NILs for QTLs of interest are constructed, and eliminates laborious hybridization work.

In this study, the NILs of three QTLs, *qGS3*, *qTGW6.2* and *qGT7*, were constructed, and effects on grain size and TGW were evaluated. The beneficial alleles of *qGS3* from GZ63-4S could increase the value by 0.21mm in GL, 0.07mm in GT, and 1.47g in TGW in homozygous NILs, which was consistent with the values of additive effect in the F_2_ and F_2:3_ population on the whole (Table 2, Table 3), suggesting that *qGS3* is a stable and pleiotropic QTL for GL, GT and TGW. *qTGW6.2* was initially detected as a QTL for TGW, but was validated to have effect on both GL and TGW in the NIL population and act in a dominant manner (Table 4). The failure in detection of *qTGW6.2* on GL in F_2_ and F_2:3_ population may be attributed to the complexity of genome background and the low variation explained, which further supported the necessity of validation of QTLs using NILs. *qGT7* was repeatedly confirmed as a QTL for GT, and had no effect on GL and GW in the F_2_, F_2:3_, and NIL populations (Table2, Table 5). Being one of the four factors of grain size, GT has received less attention, and several cloned genes conditioning GT are responsible for GL and/or GW at the same time, such as *GS2*, *GW8* [17, 44]. Therefore, *qGT7* is a good candidate for further research of the molecular mechanism underlying GT.

### Improvement of grain size, quality and yield in rice

Grain size contributes to not only grain yield, but also grain quality, especially appearance quality and milling quality [45 46]. Abundant variation of grain size is observed in rice germplasm cultivated worldwide, thus providing valuable resources for breeding or improvement of rice grain with desirable size and yield. QTLs or genes for grain size mined from germplasm resources have been or are being exploited in rice breeding programs. The *gs3* allele and *GW5* allele from 93-11, together with beneficial alleles controlling fine eating and cooking quality from Nipponbare, were introduced into the Teqing background, and the resulting lines displayed dramatic improvement in grain size and quality [47]. The *gs3* allele and the *GW7* allele from TFA were pyramided into the background of HJX74, resulting in simultaneously improvement of grain yield and quality [18]. In this study, seven QTL regions were repeated detected in both the F_2_ and F_2:3_ populations, and three of them were further validated in the NIL background, demonstrating the reality and stability these QTLs. *qGL6* and *qTGW6.2*, the two QTLs for GL located on chromosome 6, could be used in improvement of GL and KGW of GZ63-4S with the beneficial alleles from Dodda, and further in improvement of the yield and quality performance of hybrid combinations using GZ63-4S as the maternal parent. In addition, the QTLs detected in this study, together with QTLs or genes in other studies, could be combined in the breeding and improvement of rice grains with desirable size, quality and yield.

## Acknowledgement

This work was supported by grants from the National Program on R&D of Transgenic Plants (2016ZX08009003-004, 2016ZX08001002-002), and the earmarked fund for the China Agriculture Research System (CARS-01-03) of China.

## Author contributions

Ping Sun and Yuanyuan Zheng performed most of experiments and analyzed the date. Pingbo Li wrote the paper. Hong Ye and Hao Zhou constructed the F_2_ mapping population. Guanjun Gao participated in the field management. Yuqing He designed and supervised the study.

